# Simple Models For Neuroscience Research Discoveries: How Often Are These Models Used In Africa

**DOI:** 10.1101/2022.12.30.522314

**Authors:** S. K. Hamidu, U. Ahmad, R. Abdulazeez, Z. Muhammad, A.I. Alkhamis, M. Umar, A. A. Ladan, F. E. Nasr, A. Ahmad, S. Musa, J. Ya’u, W. O. Hamman, T. Yoshimatsu, S. R. Issa, M. B. Maina

## Abstract

Simple animal model systems such as *Drosophila*, Zebrafish, and *C. Elegans* have enabled numerous breakthroughs in understanding human health and disease. Their conserved biological processes, ever-expanding established procedures for handling, and amenability for molecular and genetic manipulation, in addition to the minimal ethical concerns, have made these models preferred choices in several life science disciplines globally. Owing to their cheap maintenance cost, adopting these model systems will help bridge the research gap between Africa and the Global North and contribute to advancing scientific knowledge in African universities through practical sessions. However, the extent to which these models are used across Africa is unknown. Here, we analysed the use of *Drosophila*, Zebrafish, and *C. elegans* model systems in scientific publications from African laboratories from the year 2000 to 2021. Of 1851 PubMed-indexed publications in which at least one simple animal model was mentioned, 168 used at least one of these models for the actual investigation. With an average of 21 articles per country, South Africa, Nigeria, Kenya, Egypt, Morocco, and Tunisia contributed 75% of these studies. The remaining 25% were contributed by seven other countries at 2-7 articles per country. From here, we extracted and analysed information on funding and international collaboration. This revealed that 24.4 % of the studies were exclusively funded locally, 28.57 % exclusively funded internationally, 15.5% received both local and international funding, and the rest (31.5%) were unfunded, revealing that there is satisfactory access to funds for simple animal model studies, especially from external funders. By analysing the pattern of collaborations, we show that most of these studies had international collaborations, while very few collaborated within Africa. Our work provides data on the current state of research using simple model systems in African laboratories and argues that incorporating these models will advance biomedical science research in Africa.

## INTRODUCTION

Africa is home to over a billion people with the world’s largest genetic diversity^1^. This diversity is important for understanding disease susceptibility or resistance, manifestation, and the body’s response to factors that may improve the quality of life^2^. The continent accounts for 25% of the global disease burden^3^, yet, contributes just about 2% of global scientific output^4^. As a result, Africa depends on scientific solutions or innovations from other continents, which may not necessarily be compatible with the African people. For these reasons, in addition to working on global research problems, there is a critical need for African science to also focus on developing homegrown solutions to local problems. Unfortunately, African scientists are significantly challenged for many reasons, including low funding, inadequate career development programs, lack of access to laboratory infrastructure, and supportive science policies^5,6^.

Some of the challenges affecting African scientists, including inadequate funding and lack of access to standard research facilities, can be resolved by using accessible animal models. Our recent work revealed that most African neuroscience laboratories use wild-type rodents such as mice and rats for their research^6,7^. However, these are expensive to maintain and less genetically amenable than many invertebrate and lower vertebrate model systems, and as a result, African scientists rarely use transgenic tools. Most biological processes in higher animals are largely conserved in many invertebrates^8^. Scientists have taken advantage of this to advance understanding of the physiology and pathology of human body systems. One of the most widely used organisms globally is the fruit fly *Drosophila melanogaster*, which belongs to the Drosophilidae family. It has been one of the most widely studied organisms for over a century, especially in the field of genetics^9^ neuroscience^10^, developmental biology^11^, and as a model of human diseases^12–14^.

Interestingly, many *Drosophila* species exist in the wild in Africa – their ancestral continent – and can be used easily collected and domesticated for studies^15^. Another versatile model is *Caenorhabditis elegans* (*C. elegans*). After its significance as a model was proposed in 1948^16^, *C. elegans* became popular and widely accepted by scientists globally, especially in the field of genetics and neuroscience. It shares many similar advantages with *Drosophila*^17–19^ with the additional benefit of having comparatively simpler biology, a shorter life cycle, and cheaper to maintain^20^. For vertebrate species, Zebrafish (*Danio rerio*) has become popular since its introduction in the 70s^21^. It has several advantages over other models in the field of developmental biology^22^. For example, its brain and other sense organs develop in less than 5 days in a transparent embryo. Moreover, it can be bred in large numbers within a short period. Therefore, these simple model systems, with their affordability and ever-expanding established procedures for handling and amenability to genetic manipulation ^23^, have great potential in promoting modern life sciences in Africa.

To what extent these models are used in Africa is unknown. Nonetheless, raising awareness of the advantage of using simple models will advance science in Africa and other low-resource settings. Using our recently described methodology that identifies only African-led research^6^, this study seeks to characterise the African-led biomedical research in the past 20 years to ascertain to what extent *Drosophila, C. elegans*, or Zebrafish were used as model organisms^6^. We specifically assessed the type of model organisms used in PubMed-indexed publications, their frequency, the discipline in which the research was conducted, funding sources of the research, and the availability of any local or international collaboration. Our work revealed that *Drosophila, C. elegans*, and Zebra are not widely used in African laboratories, raising the need for campaigns and funding initiatives to promote these models for basic science research in the continent.

## METHOD

### Literature Search

To assess the use of the invertebrates model in African-based laboratories, we modified our previous strategy, which involves manual data extraction of articles retrieved from PubMed ^6^. The data were procured in comma-delimited format (CSV) using the following search terms (((((((Drosophila) OR (fruit fly)) OR (fruit flies)) OR (Caenorhabditis)) OR (C. Elegans)) OR (*Caenorhabditis elegans*)) OR (zebrafish)) OR (*Danio rerio*)) AND (Name of African country). The search terms were repeated for all 54 African countries, leading to an overall 1851 articles identified and downloaded from PubMed. Unfortunately, the data does not give the basic information (the study location, funding declaration, collaboration e.t.c.) needed to achieve the objectives of this study. To obtain this information and exclude the articles that do not satisfy the inclusion criteria (Table 1), manual data extraction was done.

**Table 1.**
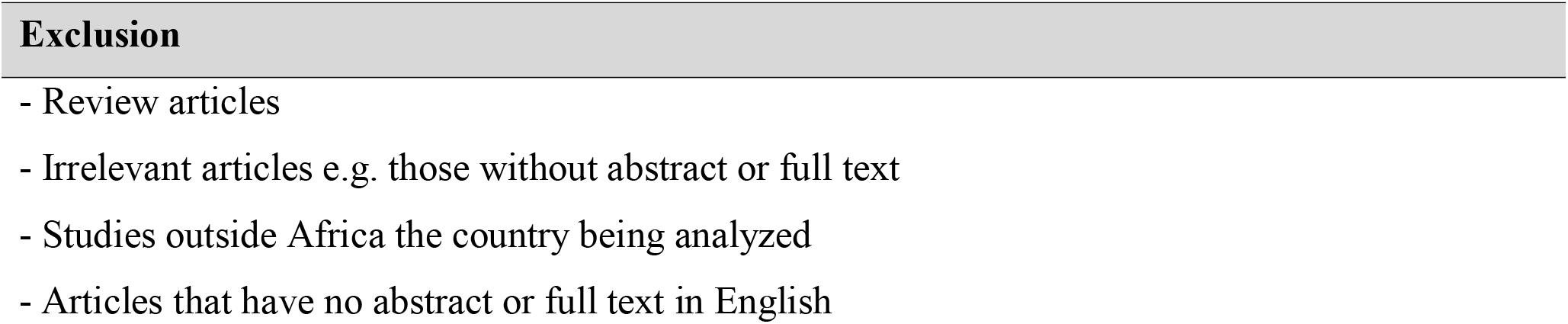
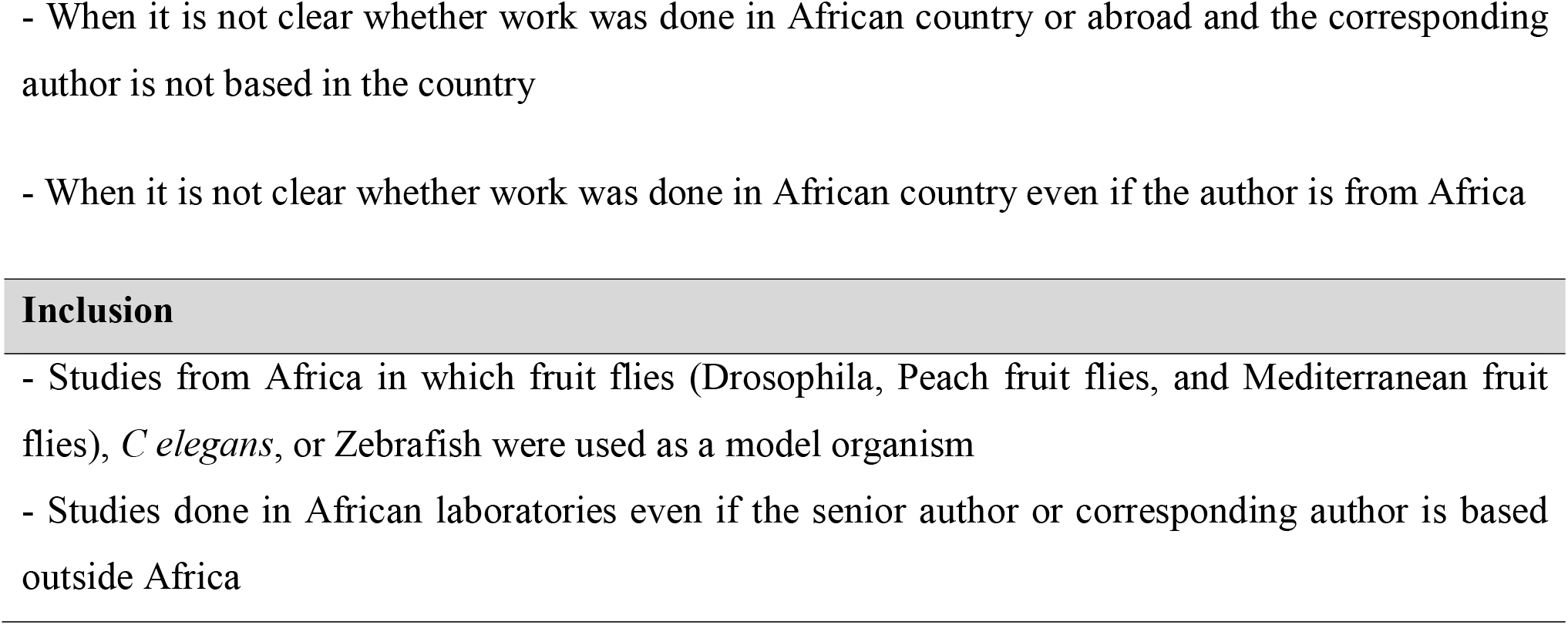
Inclusion and exclusion criteria.

### Data Availability

The data is available at https://github.com/Shkwairanga2/Simple-Animal-Model-Analysis.git

### Data Extraction

The manual data extraction was done by retrieving full texts or at least the abstracts of the articles and the details of the model organism, institutions of affiliation, and funding were extracted.

### Data Analysis

The data were analysed using Anaconda Navigator Jupyter Notebook for Python 3.8. Data exploration was done using pandas, geopandas, and the matplot library. Data visualisation was done using the Seaborn library, geoplots, and altaire packages in Python.

## RESULTS

### Number of publications between 2000 and 2020 from Africa using simple models

Out of the 1851 studies using *Drosophila, C. elegans*, or Zebrafish, whose details were downloaded from PubMed, 1683 (90.9%) articles were conducted outside Africa but at least had one author affiliated with an African institution or were articles that did not pass the specified exclusion criteria. The remaining 168 (9.10%) were conducted in African-based laboratories (Fig. 1A). An upward trend can be observed in the number of publications, especially in the last five years (Fig 1B). However, there is still a need for campaigns to encourage the adoption of simple models because the number of publications from the continent is very low (less than three (3) articles per country) compared to outputs from the Global North.

**Figure 1:**
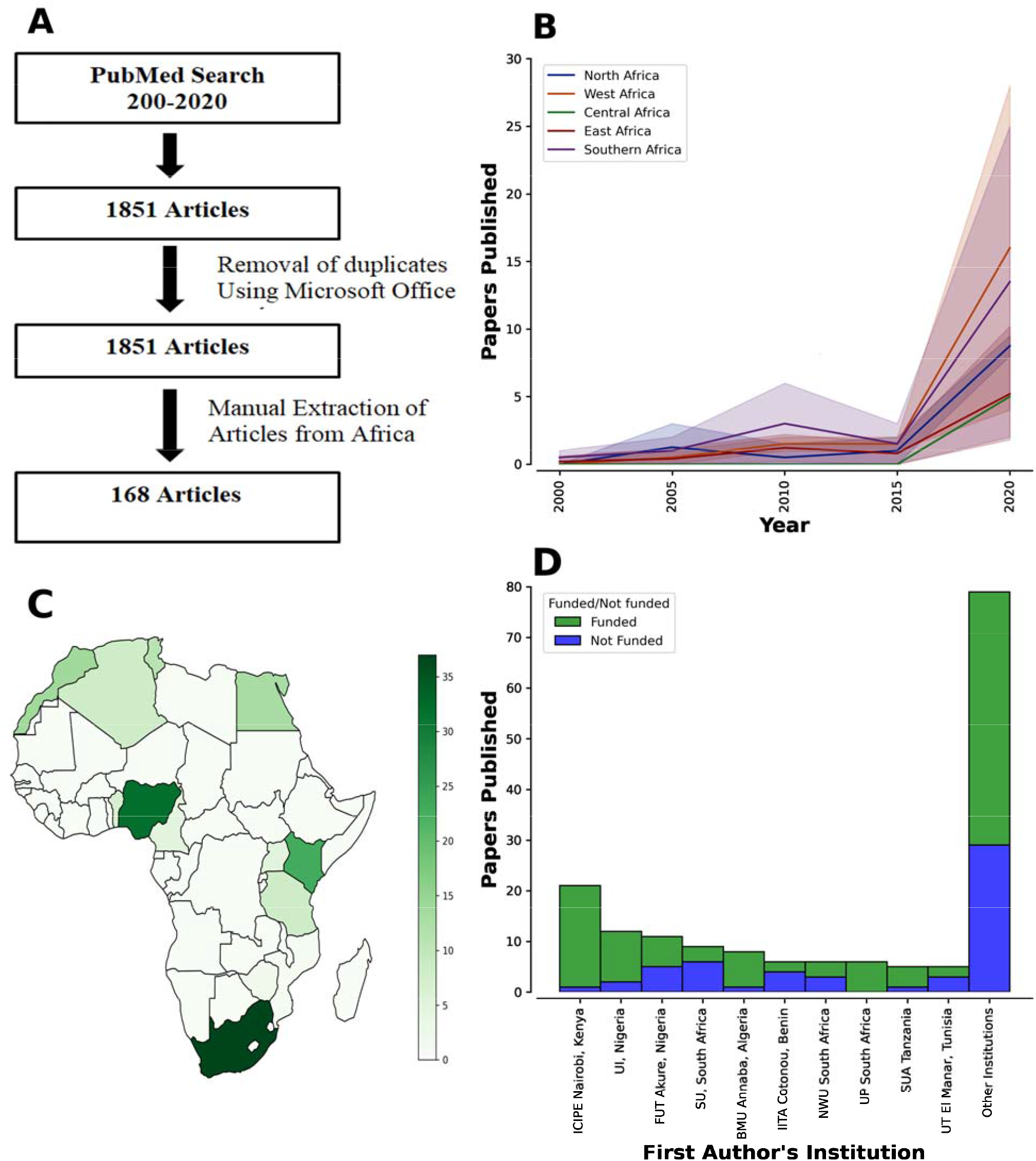
Study design and publication trends. (A) Workflow for articles retrieval and inclusion strategy. (B) Total publication per year per African region. (C) Choropleth for the number of publications by country. (D) Total number of publications by institution and their funding statuses.

75% of the studies were conducted in six (6) countries (Fig. 2). These are: South Africa (n = 37, 22.0%), Nigeria (n = 32, 19.0%), Kenya (n = 23, 13.7%), Egypt (n = 13, 7.0%), Morocco (n = 10, 7.7%), and Tunisia (n = 11, 6.5%). At 2-7 articles per country, eight other countries contributed the remaining articles: Tanzania (n = 8, 4.8%), Algeria (n = 8, 4.8%), Cameroon (n = 5, 3.0%), Benin Republic (n = 7, 4.2%), Uganda (n = 5, 2.9%), Mauritius (n = 2, 1.2%), Zimbabwe (n = 2, 1.2%) and Mozambique (n = 1, 0.6%). By region, central Africa seems to lag behind, having only six studies from Cameroon. In addition, studies from the southern part of the continent were mainly from South Africa, and most West African studies were from Nigeria.

**Figure 2:**
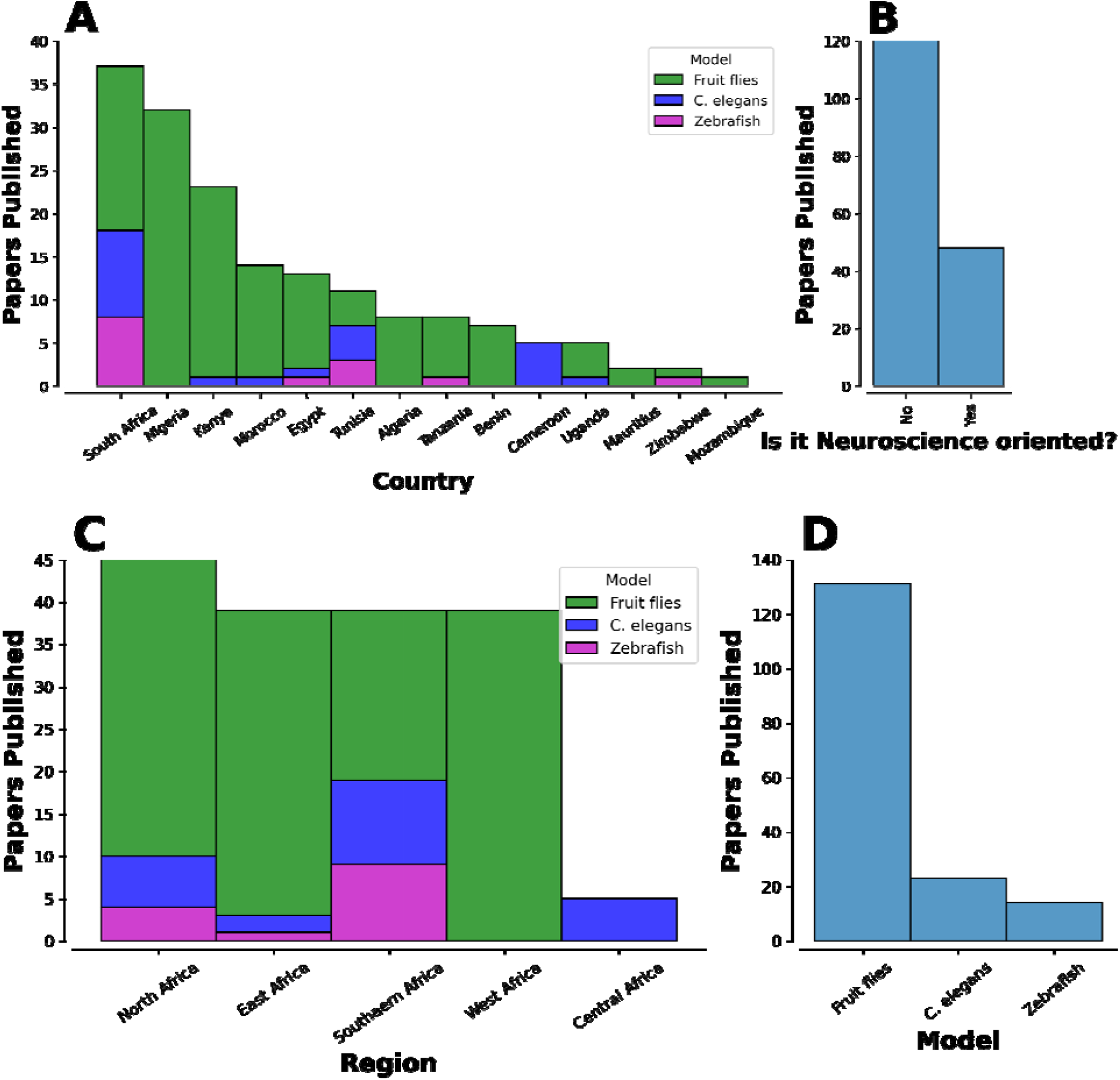
Total publications by model, region and country and neuroscience relation. (A) Total publications by country by the model organism. (B) Total publications by neuroscience relation. (C) Total publications by country by the model organism. (D) Total publications by model organism

By Institution, ten (10) institutions contributed about 53% of the studies (Fig 1D). These are: the International Centre of Insect Physiology and Ecology, Nairobi, Kenya (n = 21, 12.5%), University of Ibadan (UI), Nigeria (n = 12, 7.1%), Federal University of Technology (FUT), Akure, Nigeria (n = 21, 12.5%), Stellenbosch University (SU), South Africa (n = 9, 5.4%), Badji Mokhtar University (BMU) of Annaba, Algeria (n = 8, 4.8%), International Institute of Tropical Agriculture (IITA) Cotonou, Republic of Benin (n = 6, 3.6%), North-West University (NWU), South Africa (n = 6, 3.6%), University of Pretoria, South Africa (n = 6, 3.6%), Sokoine University of Agriculture (SUA), Tanzania (n = 5, 3.0%), and University of Tunis (UT), El Manar Tunisia (n = 5, 3.0%). Beyond this, no institution contributed up to five (5) studies. The position of the top two contributors is not surprising because ICIPE is one of the top centres for insect-based studies in Africa, and UI, Ibadan, Nigeria, has one of the main neuroscience research laboratories in Nigeria using simple models such as Drosophila.

Given that these models are mostly used for neuroscience research^24^ we categorised the articles into neuroscience and non-neuroscience-related. Surprisingly only about 32% (n=54) of the studies were neuroscience-related. Other disciplines in which these models were used are mostly in the areas of general toxicities, reproduction, pest control, morphometric studies and genetics. The most widely used model is the fruitfly (41.7%), followed by fruit flies that belong to the Tephrididae family, such as the peach fruit flies and the mediterranean fruitflies (n= 61, 36.3%). This is followed by the *C. elegans* (n= 23, 13.7%) and Zebrafish (n= 14, 8.3%).

### Research funding

Although some publishers do not require funding declaration and most African scientists do not usually acknowledge funding, 67.9% (n=114) of the studies acknowledged funding, and over 60% of the funds came from international organisations (Fig 3 A and B). The top international funders (countries) who contributed over 70% of the total funding are European countries, specifically: Germany (n=20, 30%), Italy (n=9, 13.9%), and Austria (n=5, 7.6%) (Fig 3C). By organisations, the most frequently acknowledged funders are The World Academy of Sciences (TWAS) (n=7, 10.9%), The International Atomic Energy Agency (IAEA) (n=5, 7.6%), and the German Academic Exchange Service (n=7,10.9%). By local funding, the most frequently acknowledged funders are the National Research Foundation (NRF) of South Africa (n=19, 28.8%), ICIPE, Kenya (n=7, 10.6%), and the National Fund for Scientific Research Algeria (n=6, 9.1%) (Fig 3D). It is noteworthy that all the studies from the southern part of the continent were conducted in South Africa and were locally funded, affirming the previously reported^6^ positive progress of South Africa’s policies on research development. It is also important to note that except for The IAEA, which does not fund neuroscience research, all the top funders of simple animal model studies also fund Neuroscience research.

**Figure 3:**
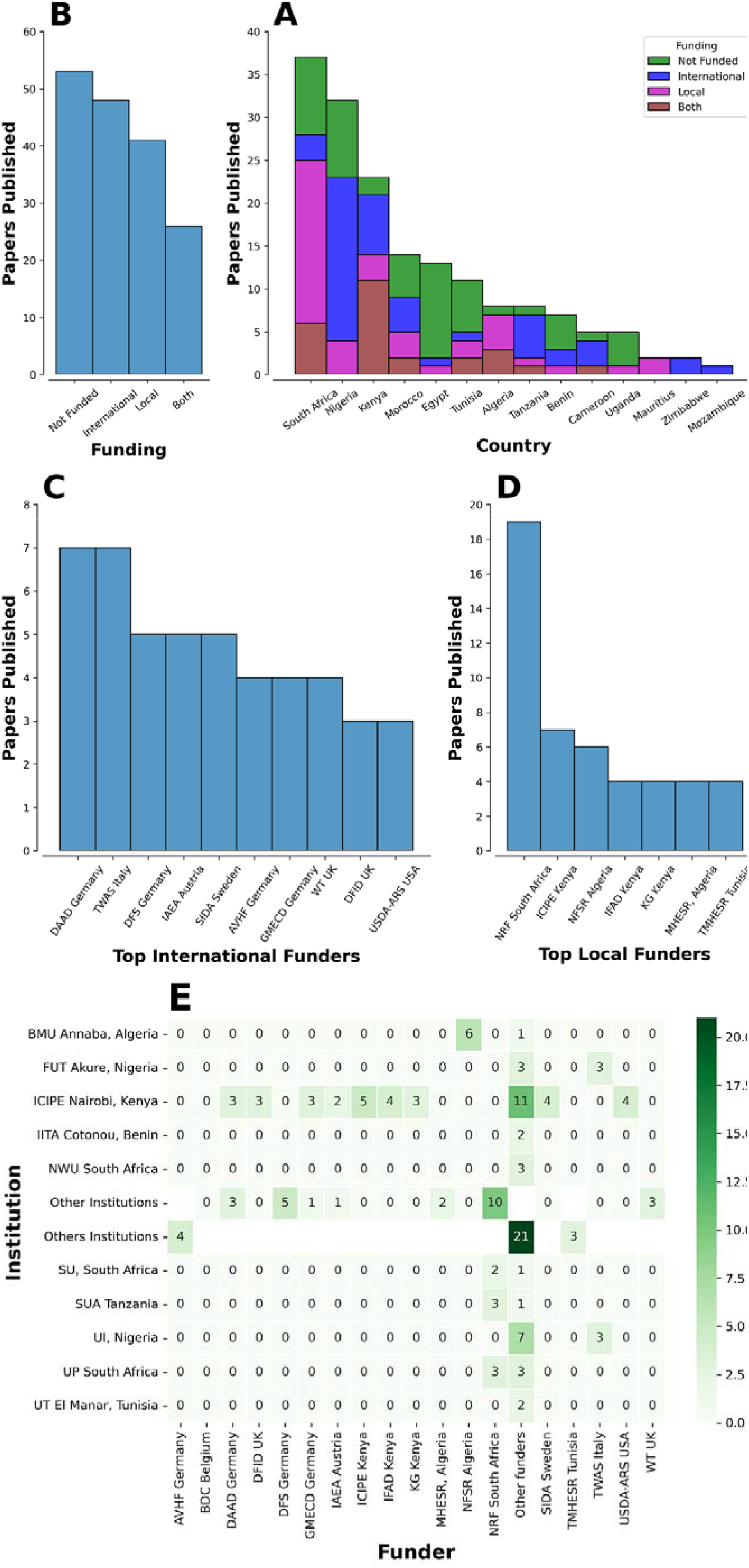
Source of funds for studies between 2000 and 2020 from Africa using the simple model. (A) Number of Articles by Funding Status. (B) Funding status by country. (C) External funders by funding agencies. (D) African funders by funding agencies. (E) Number of funding received by the top institutions from the top funders.

Funding by institutions (Fig 3E) shows that except for ICIPE and some Nigerian Institutions (UI, and FUT Akure) who also received funding from The Agricultural Research Services of the U.S. Department of Agriculture (USDA ARS USA), and TWAS, respectively, most of the top contributors (institutions) of simple animal model studies in the continent also received local funding from their respective countries and this further reveals the impact of local funding.

### Research collaboration

By international collaborations, more than half (n=87, 51.8%) of the African-based studies using the invertebrates model did not have contributions from any other country (Fig 4A). Only 13 studies (7.7%) collaborated with another African country (Fig 4B). Beyond this, all other collaborations were external, mostly with European countries. This suggests a low collaborative culture among African scientists and calls for the promotion of intra-collaboration among them.

**Figure 4:**
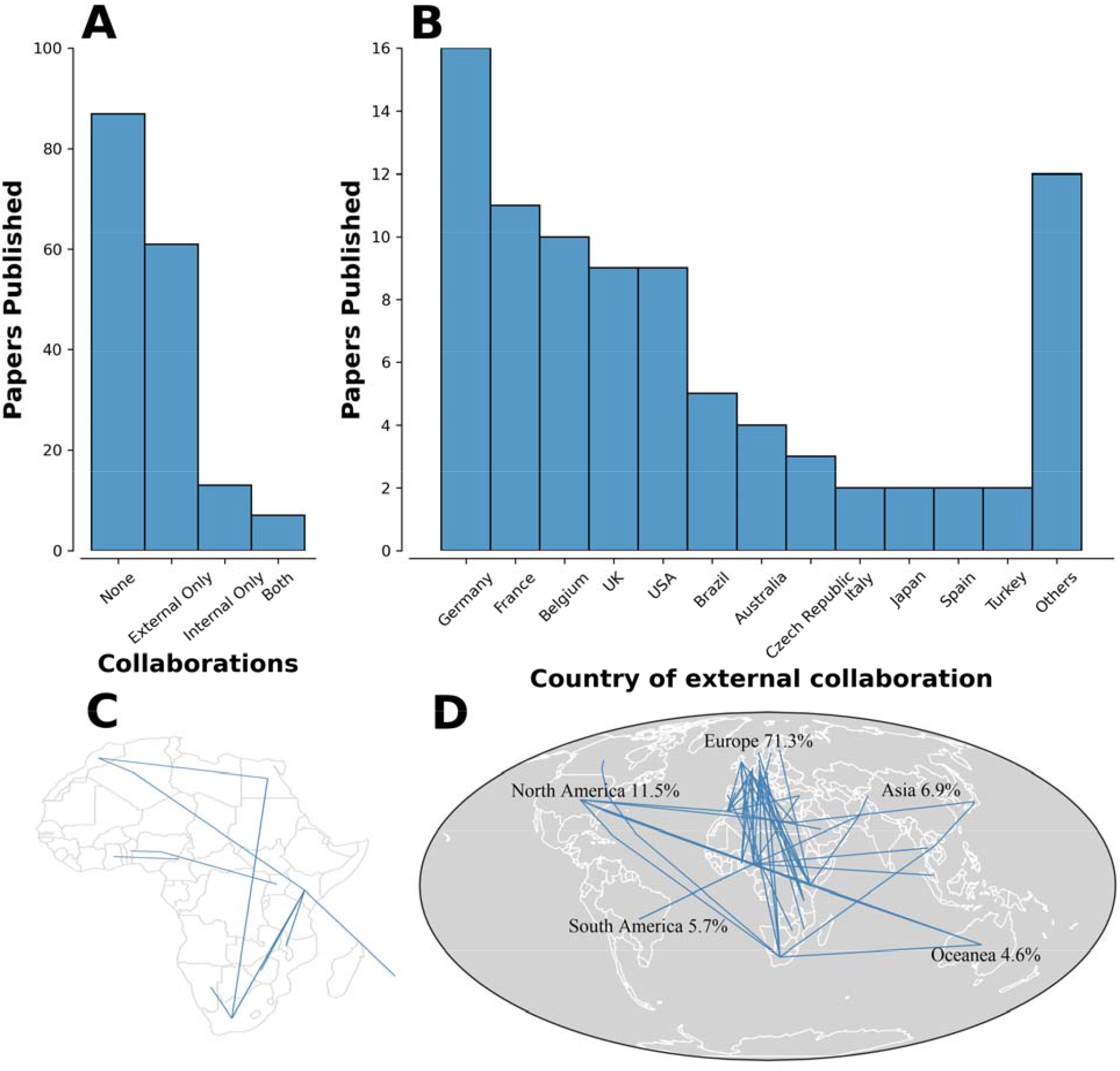
Collaborative structure for studies between 2000 and 2020 from Africa using the simple model. (A) Number of Articles by collaboration status. (B) Top external collaborators (Countries). (C) Sankey diagrams showing the intra-African collaborations. (D) Sankey diagrams showing the external collaborations.

## CONCLUSION AND DISCUSSION

The publication trend using invertebrate and lower vertebrate models agrees with the previous studies^6,25^, demonstrating that the scientific output from the African continent is mostly dominated by the six top contributors; South Africa, Nigeria, Kenya, Egypt, Morocco, and Tunisia. This is not surprising because most of these countries rank among the top economies in the continent^26,27^. South Africa has always been at the top in contributions to science because the standards of education and research in the country are higher compared to other African countries, and the South African health research policy^28^ identified and reduced the funding gap in health research by earmarking 5% of international development aid agencies project and that of program aid for the health sector to biomedical research. Nigeria’s rank as the second-highest contributor to studies using invertebrate and lower vertebrate models may also be attributable to the activities of many organizations, including TReND in Africa (www.TReNDinAfrica.org), DrosaAfrica (www.drosafrica.org) and the Neuroscience Society of Nigeria (NSN), who frequently organise workshops and awareness campaigns to foster enthusiasm in the use of simple model systems. This could be why more than 80% of the studies from Nigeria were neuroscience-related. TReND in Africa has also equipped many African scientists with skills and technology in neurogenetics, genome editing, behavioural neuroscience, and electrophysiology, among others, through workshops, donation of equipment and consumables, and the establishment of biomedical research centers^29–31^. Nigeria is one of the top beneficiaries of TReND programs. Another top contributor to African neuroscience is the International Society of Neurochemistry (ISN), International Brain Research Organisation and Dana Foundation, who provide funding support for summer schools or workshops in neuroscience or outreach programs such as brain awareness week and career talks to encourage adoption of neuroscience as the area of specialisation. By research laboratories, most of the studies from Nigeria are linked to Dr Amos Abolaji’s laboratory at the University of Ibadan, who has been leading initiatives to promote the use of Drosophila through the Drosophila Research and Training Centre (https://drosophilartc.org/), supported by many of the aforementioned organisations, among others.

The performance of Kenya, which was not ranked among the top of Africa’s strongest economies, may be due to the presence of the International Centre of Insect Physiology and Ecology (ICIPE), which is one of the major institutes for insect research in the continent. Looking at the total number of publications from the continent, it can be deduced that despite the numerous advantages of using invertebrates as model organisms, including cost-effectiveness, which can enable scientists to answer complex research questions at a low cost compared to the vertebrate models, these models are yet to be adopted widely in Africa. One of the primary reasons for the low scientific outputs from Africa is inadequate funding ^6,25,32^, and this might be because the continent has the lowest GDP per capita. Moreover, most African countries invest less than 1% of their total GDP on research development despite African Union’s recommendation. It can be observed from this study that the funding for invertebrates studies in Africa mostly comes from external sources, and this reveals the need for the scientist to engage policymakers, philanthropists, and charitable organisations in Africa to support invertebrates studies given their huge potential in advancing science in the continent. It is important to note that for the continent to establish a sustainable invertebrates research environment, local funding must be greatly improved, as can best be demonstrated by South Africa, which has a more sustainable research system than most African countries. South Africa is the only country with more than half of its simple models’ studies locally funded, and this is reflective of the fact that the country invests over 0.8% of its GDP (the highest in the continent) in research development and this could be the reason why six South African universities ranked among the top ten universities in Africa^33^.

On international collaborations, this study clearly portrays the room for improvement in intra-collaboration among African countries because an interdisciplinary approach is the best approach to efficiently address complex problems^34^. For instance, a scientist working on pest or vector control in the field of agricultural science can develop the capacity to diverge into neuroscience to be able to develop more environmentally friendly methods of vector and pest control.

Mood disorders such as depression, bipolar, and seasonal affective disorders are very common in Africa and are known to reduce the quality of life of the affected individuals significantly. These disorders may be linked to Africa’s physical and social environmental peculiarities. Unfortunately, Africa’s research capacity to understand the molecular mechanism underlying these disorders is very poor. Luckily, these cheap models are available for most neurological disorders, such as fly models of depression, dementia, and epilepsy. Also, scientists envisage an emerging opportunity for advancing molecular psychiatry through the convergence of human psychiatric genetics and neurogenetics using these models.

Taken together, it is clear that simple models are under-exploited in African labs, despite the huge potential they offer. There is a clear need for increased investment into training in the use of simple models derived from Africa or elsewhere in tackling basic research problems. While international collaborations and funding are valuable, funding and collaboration from within Africa are critical for African research’s broader impact and sustainability. Given the obvious benefits of using simple models in Africa, local and international funders need to sustain their support for seminars to increase awareness of using these models and training workshops to provide hands-on training to African scientists in using them. Importantly and owing to their low cost, funders need to provide funds to allow African scientists to establish their laboratory infrastructure, which will be key to establishing a sustainable local research environment.

## ACKNOWLEDGEMENTS

Funding was provided by the Yobe State Government, Yobe State University, Alzheimer’s Association (AARFD-22-923450), TReND in Africa, Wellcome Trust (Ref: 2020_1601_NMH) and Chan Zuckerberg Initiative (2021-240253) to MBM.

